# Capture of Viruses and Microorganisms in Aerosols Using A Newly Designed Collection System: A Proof-of-concept Study

**DOI:** 10.1101/2021.06.05.447210

**Authors:** Dapeng Chen, Alese P Devin, Emily R. Caton, Wayne A Bryden, Michael McLoughlin

**Affiliations:** Zeteo Tech, Inc., Sykesville, Maryland, United States of America

## Abstract

Aerosols contain human pathogens that cause public health disasters such as tuberculosis (TB) and the ongoing COVID-19 pandemic. The current technologies for the collection of viruses and microorganisms in aerosols face critical limitations, necessitating the development of a new type of sampling system to advance the capture technology. Herein, we presented a new type of collection system, which exploits the affinity between carbon chains and organic molecules on the surfaces of viruses and microorganisms. We demonstrated that the physical capture efficiency of the collection system was over 99% for particle sizes from 0.3 to 10 µm. We further evaluated the biochemical capture efficiency of the collection system using mass spectrometry approaches and showed that the biochemical information of viruses and microorganisms was well preserved. Coupled with well-established molecular technologies, this new type of capture technology can be used for the investigation of aerosol-related disease transmission models, as well as the development of innovative screening and monitoring tools for human diseases and public health issues.

## INTRODUCTION

Particle-containing aerosols are small-size liquid or solid particles suspended in air. Mounting evidence suggests that viruses and microorganisms are present in aerosols and play a notorious role in airborne disease transmission, including tuberculosis (TB), seasonal influenza, and COVID-19 pandemic.^1-5^ Therefore, understanding the biological role of aerosol particles is imperative to the development of prevention and treatment strategies for public health agencies.

Physical properties of aerosol particles are explored for the development of aerosol collection devices, which includes the most widely used types such as impactors, impingers, and membrane filters.^6-10^ In impactor and cyclone-type devices, particles in the air are maneuvered into an aerosol beam, accelerated through several stages of impactors equipped with nozzles decreasing in size sequentially, and deposited onto a collection surface.^11^ The analytes are extracted from the collection surface and analyzed with different molecular techniques.^11^ In impinger-type samplers, aerosol particles are directed into a liquid solution, and the collection is accomplished via the diffusion between air bubbles and the liquid medium.^9^ Since collected viruses and microorganisms are maintained in the liquid medium, cell culture assays can be conducted directly without extra extraction steps.^12^ Although impactors and liquid impingers are the most widely used devices for the collection of virus and microorganism-containing aerosol particles, they are disadvantageous due to their incomplete capture of particles of all sizes.^12^ For example, the cut-off size for impactors, identified by the resolution of nozzles, is the major challenge for the efficient collection of particles of very small diameter (<1 μm), and re-aerosolization during diffusion in impingers causes significant virus losses as flow rate increases.^12^ Fibrous filter-based technologies, such as electrets and Teflon filters, have been used for the development of portable aerosol collection systems.^10^ However, it is concluded that environmental humidity directly causes the pressure drop, as liquid or biological materials accumulate on fibrous filters, resulting in a dramatic efficiency reduction overtime. In addition, releasing of captured materials by the filter-based methds requires rigid extraction protocols that causes protein degradations. ^10^ Given these limitations, a sampling device that can collect aerosol particles in a comprehensive and complete fashion is needed.

The cell surface of a bacterium, independent of the type, is composed of various structures of organics such as the wax-like mycolic acid coating on *mycobacterium tuberculosis*.^13,14^ Those organic molecules are hydrophobic and show an outstanding affinity to alkyl chains maintained by hydrogen bonds in the water phase. This chemical feature has been employed in the analytical chemistry community for the development of retention apparatuses such as liquid chromatography columns and solid-phase extraction technologies.^15-17^ Nevertheless, the current sampling methods for the collection of viruses and microorganisms in aerosol particles entirely rely on physical properties. Methods exploiting chemical properties have not been fully explored. In this study, a carbon-chain-based system was developed and evaluated for the collection of viruses and microorganisms in aerosol particles. The collection system was applied to representative organisms, and the physical and biochemical collection efficiencies were evaluated using mass spectrometry approaches.

## EXPERIMENTAL SECTION

### Materials

All LC-grade solvents and chemicals were acquired commercially from Fisher Chemical (Waltham, MA). C18 resin beads (ID 12-20 µm) were purchased from Hamilton (Reno, NV). The 10 µm pore-size filter discs and the column sets were acquired commercially (Boca Scientific, Dedham, MA). *Escherichia coli* bacteriophage MS2 was purchased from ATCC (Manassas, VA). *Escherichia* coli K12, *Pseudomonas fluorescens* 1013, and *Yersinia rohdei* CDC 3022-85 were acquired from the Johns Hopkins University Applied Physics Laboratory. Virus titer and bacterium concentration information is provided in the online methods (Table S1).

### Aerosol generation and collection

The simulation of aerosol generation and collection was presented in Figure 1. Briefly, viruses and microorganisms were prepared in MS-grade water. The samples were loaded into an industrial Sono-Tek ultrasonic nozzle (Milton, NY; online method) to generated virus and microorganism-containing aerosols into a releasing chamber, a 50-mL conical tube. For the aerosol collection, a column (0.6 mm ID) packed with 25 mg of C18 resin beads (Figure 1a) was installed on the bottom of the releasing chamber and pulled by as diaphragm pump in a flow rate of 500 mL/min. The filters and C18 resin beads used in the current study were carefully evaluated to avoid significant pressure drop (Supporting information). A portable laser air particle counter (MetOne Instruments, Grants Pass, OR) was used to measure particle sizes from 0.3 to 10 µm. To avoid extra water vapors fusing into the diaphragm pump, a condensate collection cup, incubated in ice water, was installed after the collection column. When the collection was complete, the column was carefully removed from the system and washed with 400 µL of water three times. The wash samples were saved. The final collection products were eluted with 200 µL of 70% isopropanol into a 1.5-mL microcentrifugation tube. Samples were saved for mass spectrometric analysis.

**Figure 1.**
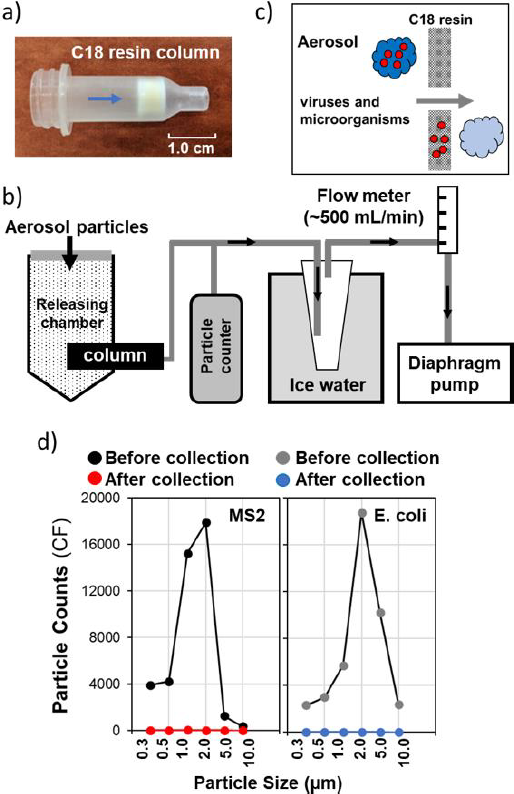
Schematic representation of the collection system that includes a C18 resin-packed column (a), the collection system components (b), the capture mechanism (c), and evaluation of the physical capture efficiency of the collection system using MS2 and E. coli (d).

### Mass spectrometric analysis of viruses and microorganisms

The protocols for the whole-cell MAIDL-TOF mass spectrometric analysis of the virus and bacteria have been well established and described in previous studies and the online methods.^18-20^ MALDI-TOF mass spectra were acquired using a Shimadzu Axima CFR-*plus* mass spectrometer in the linear mode from 1000 to 15,000 m/z. For direct infusion and nanoflow LC mass spectrometry, a LTQ orbitrap system coupling with an EASY-nLC 1000 system was used (Thermo Fisher Scientific). The flow rate for direct infusion was set to 3 µL/min. For LC-MS analysis, samples were injected into a microflow C18 column (Acclaim™ PepMap™ 100, 75 µm x 2 µm x 250 mm, Thermo Fisher Scientific) and proteins were separated using a gradient of solvent B (99% acetonitrile with 0.1% formic acid) from 5 to 65% in 90 minutes using an EASY-nLC 1000 system (Thermo Fisher Scientific). Ion fragmentation was conducted using collision-induced dissociation (CID). To improve ion fragmentation coverage, a staged-CID approach was used, where the top-down mass spectra were acquired using collision energies of 0, 10, 15, 20, 25, 30, and 35%, respectively. Monoisotopic masses were deconvoluted using Xcalibur software 3.0 (Thermo Fisher Scientific) and the fragmentation ions were interpreted using ProSight Lite (northwestern.edu).^20,21^ For richer details on top-down mass spectrometric data analysis, the reader may refer to our previous studies and the supplemental information.

## RESULTS AND DISCUSSION

### Evaluation of the physical capture efficiency on virus and microorganism-containing aerosol particles

Aerosol particles have a wide range of size distribution from sub-micro to hundreds of microns.^23^ As such, a collection system ought to be advanced enough to capture particles of all sizes in a comprehensive fashion. To evaluate the physical capture efficiency of our collection system, we generated aerosol particles that contained viruses and microorganisms ranging from 0.3 to 10 µm and measured the particle counts before and after the collection. The particle counts for MS2-containing aerosol before using the collection system were 3921, 4236, 15232, 17892, 1250, and 365 cubic feet (ft^3^, CF) at the particle sizes of 0.3, 0.5, 1.0, 2.0, 5.0, and 10.0 µm, respectively (Figure 1d, dark dots; Table S2). After the collection, the particle counts dropped to 12, 10, 52, 7, 2, and 0 CF from 0.3 to 10.0 µm, suggesting that >99% MS2-containing aerosol particles were captured (Figure 1d, red dots; Table S2). The same high-capture efficiency was also observed using E. coli-containing aerosol particles (Figure 1d, grey and blue dots; Table S2). Previous studies have shown that the capture of very small-size particles (<1 µm) is an outstanding challenge for current aerosol sampling technologies. Consequently, achieving a rigorous capture of particles from 0.3 to 10.0 µm underscores a major advantage of our collection system^12^. Several factors contribute to the excellent capture efficiency of this collection system. The packed C18 resin beads are 12-20 µm in diameter, resulting in 0.3 µm to 25 µm in pose size of the column, depending on the shape of the beads. The length of the packed beads is around 3 mm, and this design provides a longer contacting path and expands the contact areas to much larger capacities than filter membrane-based collection systems which has a thickness less than 0.1 mm (us.vwr.com). ^23^ This physical characteristic of the column provides a comprehensive coverage of all size aerosol particles, contributing to a complete capture.

### Capture of MS2 and bacteria using the collection system revealed by mass spectrometry

Three types of bacteria were aerosolized to generate particles and characterized using MALDI-TOF mass spectrometry after the collection (Figure 2). The results showed that reported bacterium signatures were well observed in elution samples with MALDI-TOF mass spectrometry (Figure 2). ^24-27^ To rule out the possibility of carry-over and cross-contamination, samples collected from the wash steps were evaluated and no bacterium signatures were observed in those samples (Figure 2, middle images). In fact, after a quick centrifugation on the elution samples, bacterium materials can be visualized as pale pellets on the bottom of the tube, which highlights the exceedingly strong capture capacity of the collection system (Figure S1). In addition, the signal-to-noise ratios between the control and elution samples were indistinguishable in all three bacterium types, suggesting that the captures were complete and all bio-materials were well-preserved in the collection (Figure 2). By using three types of bacteria, we demonstrated that our collection system has the capacity to capture microorganisms in aerosol particles in an unexpurgated fashion.

**Figure 2.**
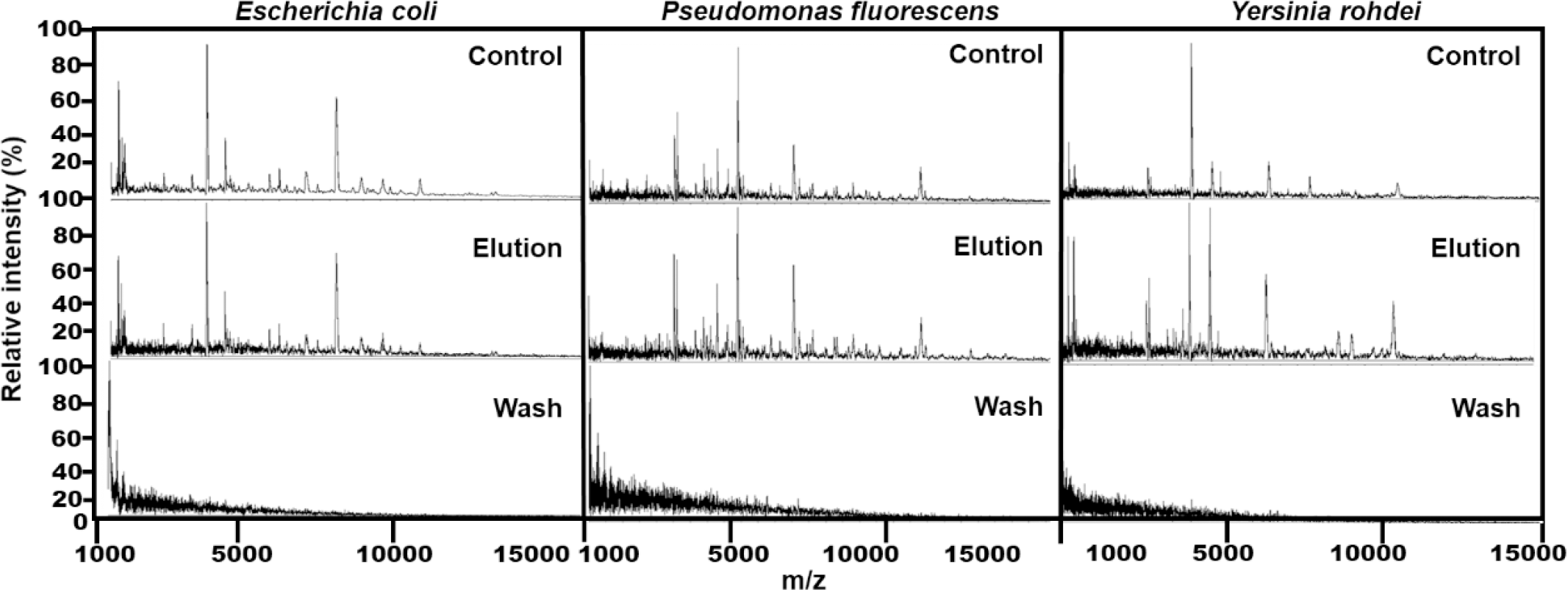
Characterization of bacteria in control samples and samples collected from elution and wash steps using MALDI-TOF mass spectrometry.

The COVID-19 pandemic is caused by the spread of SARS-CoV-2. Certain non-physical contact transmission cases suggest the virus may be contained in air aerosols and travel for a long distance to cause the infection.^28^ Therefore, it is of great importance to develop a device that has the capacity to capture viruses in aerosol particles. Since most viruses contain a protein shell, granting the certainty that our collection system ought to be able to thoroughly capture capacity to capture viruses thoroughly.^18^ In our study, MS2 was used as a representative virus model to evaluate the capture capacity of our collection system. It should be noted that MS2 is a small size virus (∼27 nm) and thus more physically challenging to capture than larger viruses such as severe acute respiratory syndrome coronavirus and influenza viruses. MALDI-TOF mass spectrometric analysis showed that the historical biomarker of MS2, capsid protein, was observed in the elution sample after the collection (Figure 3a). ^18^ The identity of MS2 capsid protein was unanimously confirmed using a top-down mass spectrometry approach in which high-confidence statistical scores were constructed by matching the experimental fragmentation ions to *in silico* protein fragmentation patterns (Figure 3b). Whole-cell MALDI-TOF mass spectrometry characterization and top-down protein identification confirmed that our collection system has the capacity to capture viruses in aerosol particles.

**Figure 3.**
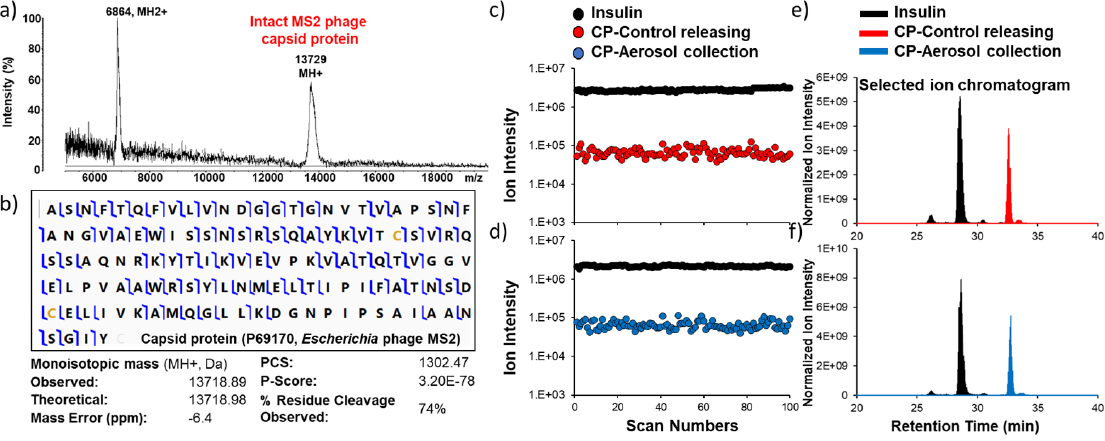
Characterization of MS2 using MALDI-TOF mass spectrometry (a) and identification of MS2 capsid protein using top-down mass spectrometry (b). Direct infusion mass spectrometric analysis of MS2 capsid protein (CP) in control samples (c) and samples collected from aerosol particles (d). Nanoflow-LC mass spectrometric analysis of MS2 capsid protein (CP) in control samples (e) and samples collected from aerosol particles (f).

### Evaluation of the biochemical capture efficiency on viruses and microorganisms in aerosol particles

The success of using the collection system to capture three types of bacteria and the virus in aerosol particles motivated the investigation of the biochemical capture efficiency in a more quantitative way. For this purpose, targeted ion intensities generated in two sample introduction methods in top-down mass spectrometry, direct infusion and nanoflow LC mass spectrometry were used for the quantitative analysis. Direct infusion mass spectrometric analysis showed that the signal intensity of the referee insulin was around 3.6E6 (Figures 3c and d, dark dots). The signal intensity of MS2 capsid protein was around 1.1E5 in both control and capture samples, suggesting no obvious sample loss during the collection (Figures 3c and d, red and blue dots). This observation was further confirmed in the nano-flow LC mass spectrometry, the gold standard for molecule quantitative analysis. The results showed that the peak areas and ion intensities of MS2 capsid protein were identical between the control and capture samples, suggesting that the collection system preserved a high biochemical capture efficiency (Figures 3e and f). Several factors could contribute to the superior capture efficiency of our collection system. It is well known that C18 materials show great retention capacity to organic molecules as 25 mg of C18 resin beads used in our study could support as much as 2.5 mg of organic materials.^29^ As previously described, the thickness of the packed C18 resin beads allows a longer retention time and thus guarantees complete contact between carbon chains and organic molecules on the surfaces of viruses. In addition, aerosol particles and the contained viruses and microorganisms were directly exposed to the capture materials, which minimized the “wall-losses” effect that is present in other aerosol technologies where particles are deposited on the side of the device rather than being collected.^12^ Our quantitative analysis on MS2 capsid protein demonstrated that the biochemical capture efficiency of the collection system was outstanding.

## CONCLUSIONS

In this study we presented a new type of collection system for the high-efficiency capture of viruses and microorganisms in aerosol particles. The collection system has the capacity to collect aerosol particles of all sizes while preserving biochemical information. This collection system is simple and thus gives fresh impetus for the development of a rapid point-of-care device for the collection of viruses contained in human breath and aerosols, particularly aimed at seasonal flu infection and the current COVID-19 pandemic. In this study, we demonstrated the effectiveness of the collection system using mass spectrometry approaches for the identification of viruses and bacteria in aerosol particles. As expected, other molecular technologies, such as nucleic acid amplification and immunoassay, can be feasibly applied to the collection products for their identification.^30,31^ In addition, since no vacuum pump system is applied in this collection system, it can be treated as a “soft technique” and thus the viability of viruses and microorganisms can be preserved, suitable for microbiological culture studies.^12^ Human exhaled air produced from coughing or sneezing contains plenty of biological materials.^32-34^ Therefore, the rich analysis of these biological materials is not limited by analytical technologies but rather by a lack of an effective collection method.^12, 35^ Considering the advantages of our collection system, this new type of collection system has a wide range of applications, such as the investigation of airborne disease transmission mechanisms and the development of diagnostic and monitoring tools for human diseases and environmental hazards.

## Supporting information

STable01

## ASSOCIATED CONTENT

The Supporting Information is supported with the communication of the authors.

## AUTHOR INFORMATION

### Corresponding author

*To whom correspondence should be addressed: Dr. Dapeng Chen

### Author contributions

D.C, M.M, and W.A.B designed the experiments. D.C. collected aerosol particle samples. D.C. E.R.C, and A.P.D conducted mass spectrometry analysis and other analysis. D.C. drafted the main manuscript. All authors understood and agreed the results. All authors approved the manuscript.

### Notes

Dr. Dapeng Chen, Ms. Emily Caton, and Ms. Alese Devin are employees of Zeteo Tech, Inc. Dr. Wayne A Bryden is the President and CEO of Zeteo Tech, Inc. Mr. Michael McLoughlin is the Vice President of Research of Zeteo Tech, Inc. This subject matter of this paper was previously disclosed in a pending and unpublished U.S. Provisional Patent Application assigned to Zeteo Tech, Inc.

## ACKNOWLEDGEMENTS

We thank the Johns Hopkins University Applied Physics Laboratory for providing all bacterium samples.

