## Supplementary material for "Capture of Viruses and Microorganisms in Aerosols Using A Newly Designed Collection System: A Proof-of-concept Study": STable01

#### A Novel System for the Capture of Viruses and Microorganisms in Aerosols

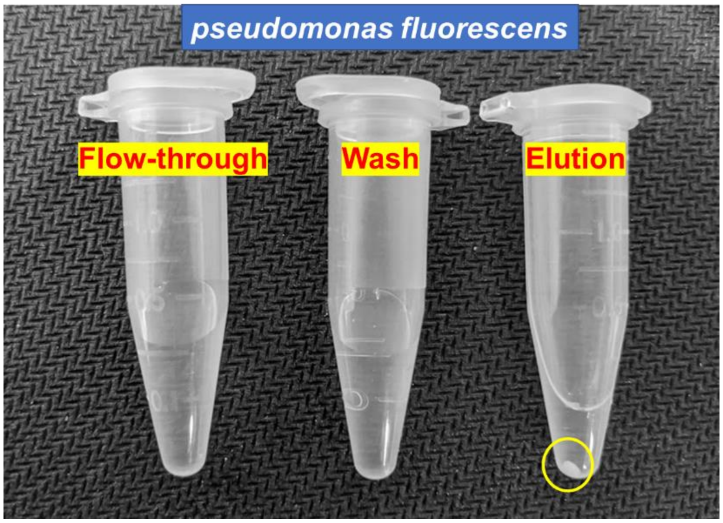


Figure S1. Visualization of bacterial materials after capture using the collection system in flow-through, washed, and eluted samples. The bacterial material was visualized as a pale pellet and indicated in the yellow circle in eluted samples. Such pale pellet was not shown in either flow-through or washed samples.

### Methods

**Release of viruses and microorganisms-containing aerosol particles.** Generally, 20 µL of materials were added in the SonoTek nozzle for aerosol generation. To avoid sample accumulation in the nozzle, the nozzle was washed twice with 200 µL of water consistently and the aerosol collection was conducted for 10 minutes to achieve a complete sample releasing and collection.

**MALDI-TOF mass spectrometry.** Collected samples were applied on the MALDI plate. Then, 2 µL of CHCA (10 mg/mL) prepared in 70 acetonitrile and 0.1% TFA was added to the sample. After the samples dried completely, the plate was inserted into the MALDI-TOF mass spectrometer. MALDI-TOF mass spectra were obtained in linear mode with a 337 nm N2 laser (laser power: 92) and all spectra were collected as an average of 1000 profiles from 1000-15000 m/z.

**Direct infusion mass spectrometry.** Samples were introduced to an LTQ orbitrap mass spectrometer via electrospray ionization by direct infusion (flow rate: 3 µL/min). Positive ion calibration solutions (Thermo Fisher Scientific) were used for ion source optimization and mass accuracy test.  For initial data acquisition, a resolving power of 60, 000 at 200 m/z was used for the positive ion mode in the mass range of 400 – 2000 m/z.

**Nanoflow LC mass spectrometry.** Briefly, protein samples were separated using a high-performance EASY nLC 1000 liquid chromatograph system (Thermo Fisher). The samples was desalted and concentrated in an Acclaim PepMap trap (2 µm × 20 mm) and subsequently separated on an Acclaim Pepmap C18 column (75 µm × 25 cm x 2 µm) at a flow rate of 0.3 µL/min using a linear gradient of 5−55% solvent B (99% acetonitrile and 0.1% formic acid) for 90 min. The column oven and autosampler temperatures were set to room temperature. Mass spectra were acquired on an LTQ orbitrap mass spectrometer (Thermo Fisher). A resolving power of 60000 was used to acquire both the precursor and fragment ions. The automatic-gain-control (AGC) target was defined as 3E6 ions during both the precursor- and fragmentation-ion acquisitions. All MS/MS spectra were produced in the data-dependent mode using high-energycollision-induced dissociations (CID) with 0-35 % collision energy.

**Data analysis for top-down mass spectrometry.** Top-down mass spectrometric identification of the MS2 capsid protein was conducted using the ProSight Lite software. Product-ion spectra were deconvoluted using the Xtract protein in the

Xcalibur 3.1 Qual Browser (Thermo Fisher). CID generates b and y ions so as the fragmentation method was set to “CID”. Mass tolerance was defined as 10 ppm. Protein sequence of MS2 capsid protein was extracted from uniprot.com.

**Effects of C18 bead size, bedding quantity, and inlet filter pore size on pressure drop of the collection system.** Membrane-based aerosol collection devices are known to have clogging and pressure drop issues, which is caused the accumulation of water droplets and environmental particles during the collection. The effects of C18 bead size, bedding quantity, and inlet filter pore size were evaluated in this study. For this purpose, the flow rate of the diaphragm pump was set to 2.5 L/min and the flow rate was monitored for 30 min during the collection. The pressure drop testing was performed using either HPLC-grade water or E. Coli vegetative cells.

Two types of C18 beads with 10 and 20 µm sizes were tested (Figure 1). The results showed that the flow rate of 20 µm C18 beads was ~750 mL/min when using HPLC-grade water and ~600 mL/min using E. Coli. The flow rate significantly dropped when using 10 µm C18 beads. The flow rate of 10 µm C18 beads dropped from ~250 mL/min to 50 mL/min during the 30 min collection. The pressure drop worsened when using E. Coli vegetative cells. The flow rate dropped from ~100 mL/min to 3 mL/min in about 5 min. The results clearly showed that pressure drop caused by water droplets and pathogen particles was an outstanding issue when using C18 beads with small sizes. Based on our results, the size of the C18 beads used for building an aerosol collection system should be larger than 20 µm.

The effects of C18 bead quantity and inlet filter size were evaluated (Figure 2). The results showed that the increase in bedding quantity would significantly lower the flow rate. When using 40 mg of 20 µm C18 beads, the flow rate dropped from ~700 mL/min to 150 mL/min when comparing to 25 mg C18 beads after 30 min of HPLC-grade water collection. Two types of inlet filters used for C18 beads packing were evaluated for the pressure drop testing. The flow rate dropped from ~600 mL/min to ~250 mL/min when comparing 35 µm pore size filter with 10 µm after 30 min of E. Coli vegetative cell collection.

The evaluation of pressure drop of C18 beads in our study suggested that several factors significantly affect the flow rate of the collection system, including the size of the beads, the quantity used, and the choose of the packing filter type. The optimization and evaluation on those factors are critical to avoid significant pressure drop and clogging issue when developing a collection system using C18 beads.

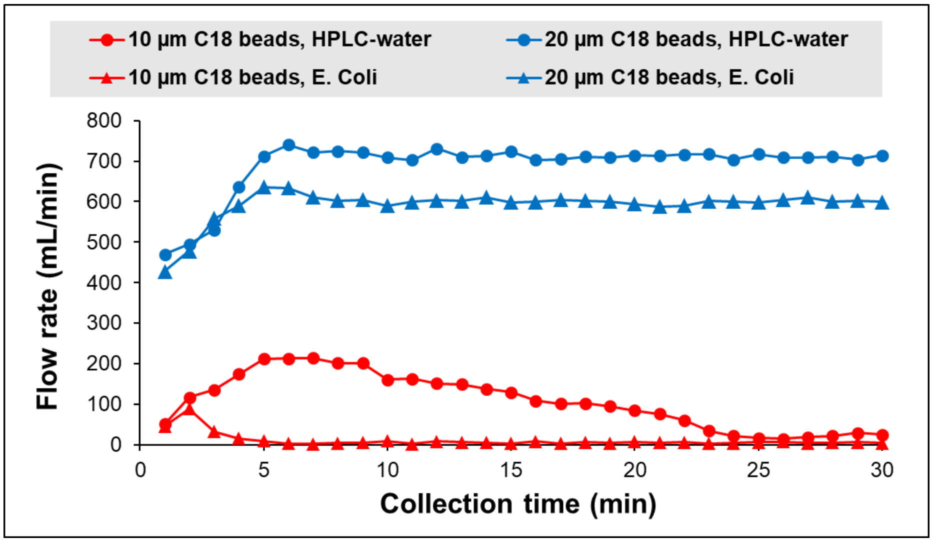


Figure S2. The effect of C18 bead size on the pressure drop of the collection system.

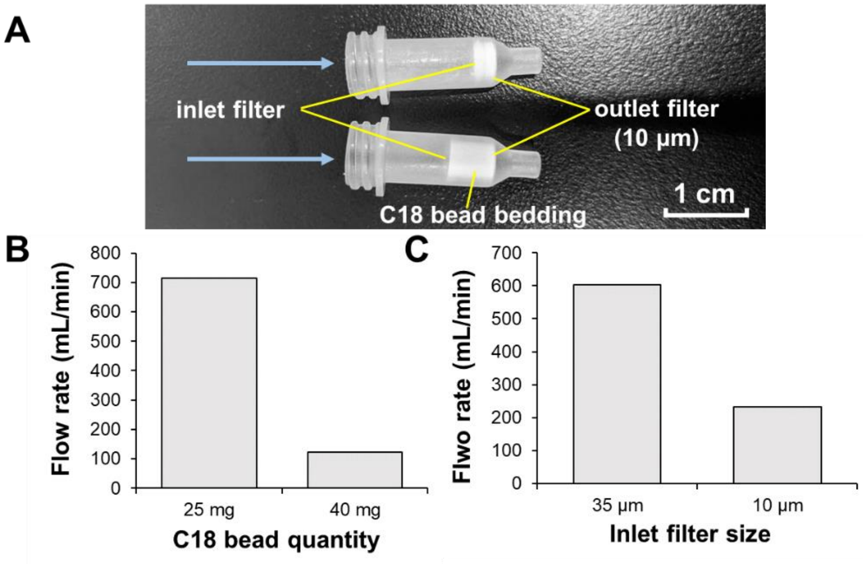


Figure S3. The effects of C18 bead bedding quantity and inlet filter pore size on the pressure drop of the collection system.

**Supplementary Tables:**

Table S1: Virus and bacterium product information.

Table S2: Capture efficiency evaluation.
